# Cheminformatics tools for analyzing and designing optimized small molecule libraries

**DOI:** 10.1101/358978

**Authors:** Nienke Moret, Nicholas A. Clark, Marc Hafner, Yuan Wang, Eugen Lounkine, Mario Medvedovic, Jinhua Wang, Nathanael Gray, Jeremy Jenkins, Peter K. Sorger

**Author notes:** Present address: Department of Bioinformatics and Computational Biology, Genentech Inc., 1 DNA Way, South San Francisco, California 94080, USA. Address correspondence to: Peter K. Sorger, cc, Harvard Medical School, Warren Alpert 444, 200 Longwood Avenue, Boston, Massachusetts 02115.

## Abstract

Libraries of highly annotated small molecules have many uses in chemical genetics, drug discovery and drug repurposing. Many such libraries have become available, but few data-driven approaches exist to compare these libraries and design new ones. In this paper, we describe such an approach that makes use of data on binding selectivity, target coverage and induced cellular phenotypes as well as chemical structure and stage of clinical development. We implement the approach as *R* software and a Web-accessible tool (http://www.smallmoleculesuite.org) that uses incomplete and often confounded public data in combination with user preferences to score and create libraries. Analysis of six kinase inhibitor libraries using our approach reveals dramatic differences among them, leading us to design a new *LSP-OptimalKinase* library that outperforms all previous collections in terms of target coverage and compact size. We also assemble a mechanism of action library that optimally covers 1852 targets of the liganded genome. Using our tools, individual research groups and companies can quickly analyze private compound collections and public libraries can be progressively improved using the latest data.

## Introduction

The size and diversity of small molecule libraries is an important design criterion: the larger the number of compounds, the greater the probability of finding a high affinity hit and the more protein targets that can be included; the smaller the library, the greater the feasibility of studying dose-response relationships, drug combinations and sensitizing conditions^1–6^. Focused small molecule libraries (typically comprising ˜ 30-3,000 compounds) are widely used to identify potential therapeutic targets, study biological processes and uncover drug repurposing opportunities^7^. Focused libraries are particularly advantageous (relative to large compound collections) with phenotypic assays, as illustrated by the identification in such collections of “first-in-class” drugs^8–10^ such as Vorinostat^11^ and Miglustat^12^ and the repurposing of drugs such as LY500307 (Erteberel) for schizophrenia.^13^ Despite this, few data-driven approaches exist to evaluate existing compound collections or design new ones.

Many focused libraries concentrate on a single druggable multigene family, such as kinases or ion-channels, FDA-approved drugs or the *liganded genome*^14^ (the subset of proteins bound by at least three compounds at K_i_ < 10 µM); libraries against the liganded genome are commonly referred to as “Mechanism of Action” (MoA) libraries because they can be helpful in dissecting biological mechanism^15^. Such libraries have become available through public initiatives^16^, outreach by pharmaceutical companies^17^, commercial small molecule vendors and expert-curated lists of “optimal” compounds^18–20^, (e.g. www.chemicalprobes.org). In principle, choosing a library should be relatively simple: optimal libraries include as many bioactive and highly selective compounds as necessary to cover a target class of interest, with the possible addition of approved and investigational therapeutics. Achieving this in practice is difficult, since many compounds exhibit polypharmacology (that is, they bind to multiple protein targets) and data on polypharmacology are frequently incomplete. It can therefore be difficult to determine whether a drug-induced phenotype of interest is due to inhibition of the nominal target, a secondary target or both^18^.

In this paper, we describe a set of algorithms and software tools for optimizing the composition of focused small molecule libraries based on binding selectivity, structural diversity, similarity in cell-based assay activities and stage of clinical development. Our approach aims to minimize the number of compounds in a library while preserving diversity and other desirable characteristics. We compare six kinase-focused libraries and find that they vary substantially in chemical diversity and the range of targets covered. We therefore assemble a new *LSP-OptimalKinase* library with properties superior to any existing compound collection in terms of target coverage and compound selectivity. We also generate a compound list for an *LSP-MoA* library that optimally targets the liganded genome. Since libraries are created using software, they can easily be updated as new information on small molecules and their targets becomes available.

## Results

In this study, we use four types of information curated in ChEMBL and other data sources to analyze small molecule libraries: (i) chemical structure, (ii) “*target data*” (iii) “*nominal target”* and (iv) “*phenotypic data*” (from drug-treated cells). Target *“dose-response data”* are most commonly derived from enzymatic assays with recombinant or purified proteins at >5 drug concentrations, yielding curves from which K_i_ or IC_50_ values can be derived. Target *“profiling data”* primarily involves assays with large panels of recombinant proteins, or proteins present in cell-free extracts, and yields information on binding to many different targets, which is often expressed as percent activity against each target (see for example DiscoverX KINOMEscan (https://www.discoverx.com/home) and Kinativ (http://www.kinativ.com/)^21–23^ “*Nominal target”* is the target a small molecule was designed to agonize or inhibit, or that it is generally assumed to target, but it may not be the highest affinity or most biologically significant. “*Phenotypic data*” comprise experimental data from cell-based assays in which drug activity is measured using a morphological, biochemical, genetic or functional readout but drug-target engagement is not determined directly (e.g. drug-induced changes in proliferation, apoptosis, migration, immune-cell activation, transcription factor translocation, etc.).

The data sources used in this work comprised the ChEMBLV22_1 database^24^, which draws on data published in the scientific literature, patents, PubChem bioassays, FDA approval packages and other resources^25^; kinome-wide screens from the International Centre for Kinase Profiling (http://www.kinase-screen.mrc.ac.uk); kinome-wide screens from LINCS^16^; and in-house curation of nominal targets (Figure S1). To correctly combine data for a single compound under different names (e.g. OSI-774, Erlotinib and Tarceva®) we matched chemical structures using the Tanimoto similarity of Morgan2 fingerprints (hereafter *“structural similarity”*)^26^.

Our analysis of focused libraries involves six widely available inhibitor libraries with public annotations (Table S1): (i) the SelleckChem kinase library (abbreviated SK), a set of 429 kinase inhibitors assembled by the commercial supplier SelleckChem; (ii) the “Published Kinase Inhibitor Set” (PKIS), a set of 362 compounds assembled by Glaxo-Smith-Kline; (iii) the Dundee compound collection (Dundee), a set of 209 kinase inhibitors screened for biochemical activity at the University of Dundee; (iv) the EMD kinase inhibitor collection (EMD), a set of 266 kinase inhibitors sold by Tocris Bioscience; (v) the HMS-LINCS small molecule collection (LINCS), a set of 495 small molecule probes and drugs focused on kinase inhibitors that was assembled by the NIH-funded HMS-LINCS center; and (vi) SelleckChem Pfizer licensed collection (SP), a set of 94 molecules licensed by Pfizer for sale by SelleckChem (see Supplementary Table 1 for references and Web links). In aggregate these libraries contain 1417 unique compounds that were mapped onto ChEMBL identifiers (Methods; Tables S2 and S3C).

### Chemical similarity within libraries

With respect to composition, we found that the LINCS and SK libraries shared ˜50% of their compounds, making them the most similar among the six libraries examined (Figure 1A). In contrast, 350 of 362 compounds in the PKIS library were unique. To visualize structural diversity, we plotted chemical similarity against itself; the presence of highly similar compounds (analogues) gives rise to off-diagonal clusters. The LINCS library generated few such clusters whereas the PKIS library was dominated by them (Figure 1B). By way of illustration, a 3-compound cluster of LINCS analogues comprised the CDK inhibitor Seliciclib, and two close relatives, Olomoucine II and (S)-CR8 (Figure 1B bottom). Seliciclib progressed to phase II for nasopharyngeal carcinoma, but failed to meet its primary endpoint of improving progression free survival^27,28^, while the other compounds are reported to have improved cellular potency and selectivity^29,30^. A 9-compound PKIS cluster contained a set of close analogs, none of which have progressed into clinical development (Figure 1B, bottom). By scoring the frequency and sizes of clusters having structural similarity ≥0.7 we find that the LINCS and Dundee collections are the most structurally diverse, PKIS the least, and SK, SP and EMD in-between (Figure 1C). This arises because LINCS, Dundee and SK compounds were drawn from different drug discovery campaigns across multiple companies and research groups, whereas the PKIS library was derived from structure-activity studies inside a single company.

**Figure 1.**
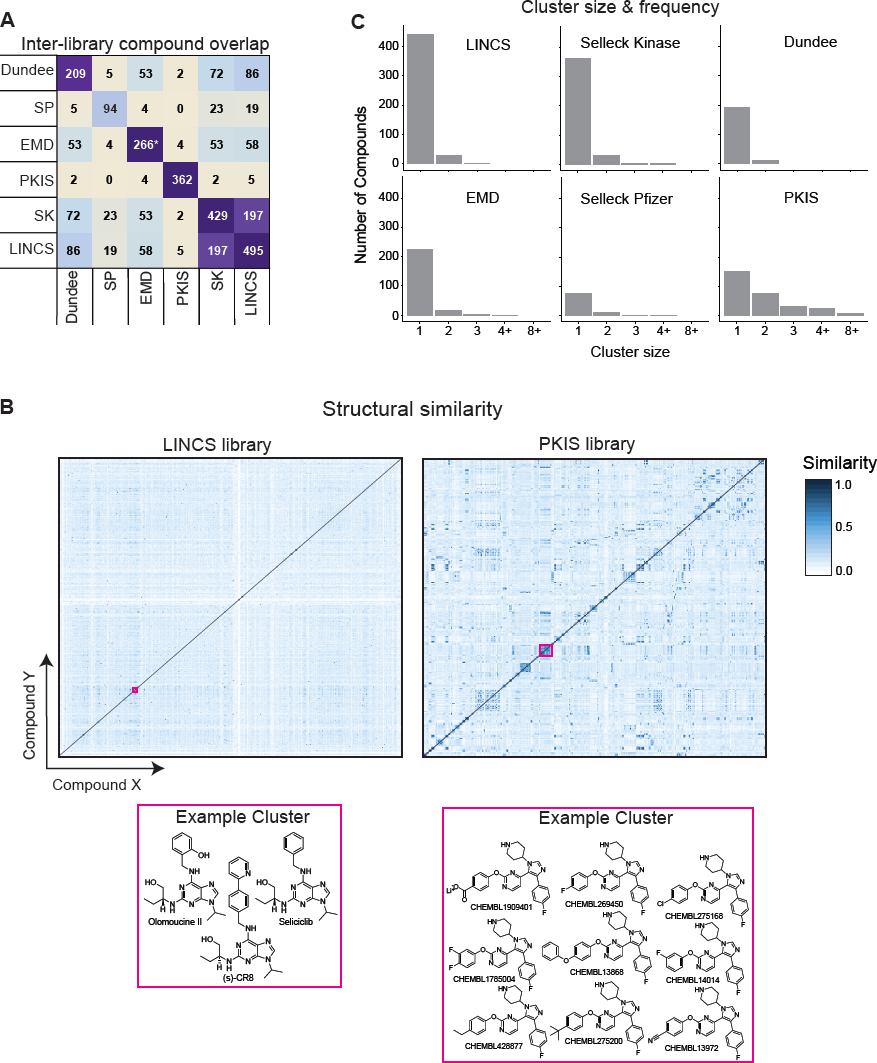
Structural comparisons across the libraries. (A) Number of identical compounds in six widely used libraries. Compounds were matched based on chemical structure; Dundee denotes a compound assembled by the University of Dundee; SP the SelleckChem Pfizer licensed collection; SK denotes the SelleckChem kinase collection; EMD a kinase inhibitor collection sold by Tocris Bioscience; PKIS the “Published Kinase Inhibitor Set” assembled by Glaxo; and LINCS, the HMS-LINCS small molecule collection (see Table S1 for references and Web links). Asterisk highlights library files provided by EMD that did not contain the same number of compounds as advertised online. (B) Structural similarity for LINCS and PKIS libraries calculated using the Tanimoto similarity of Morgan fingerprints (at 2Å); molecules in two representative clusters are shown below. Data on these compounds could not be retrieved through customer service; thus only structures present in the available library files are considered in this paper. (C) The distribution of compound clusters for different libraries (with a similarity cut-off of 0.7).

### Phenotypic similarity and clinical development

To assess the diversity of phenotypes elicited by different compounds, we constructed phenotypic fingerprint (PFP) vectors that quantify activity in a wide range of phenotypic assays curated in ChEMBL. The length of PFP vectors ranged from 5-58 entries (median = 11) for pre-clinical molecules and investigational drugs and 20-870 assays (median =440) for approved drugs. For example, the vectors for Imatinib and Nilotinib had 303 assays in common and yielded a Pearson correlation of r=0.82, whereas Imatinib and Dasatinib had 426 assays in common and yielded a Pearson correlation of r=0.16 (Figure 2A). These correlation values mirror with binding conformations that the drugs assume with the BCR-ABL oncogene: Imatinib and Nilotinib are type II inhibitors, locking ABL in its inactive conformation^31^, while Dasatinib is a type I inhibitor and targets ABL in its active conformation^32,33^.

**Figure 2.**
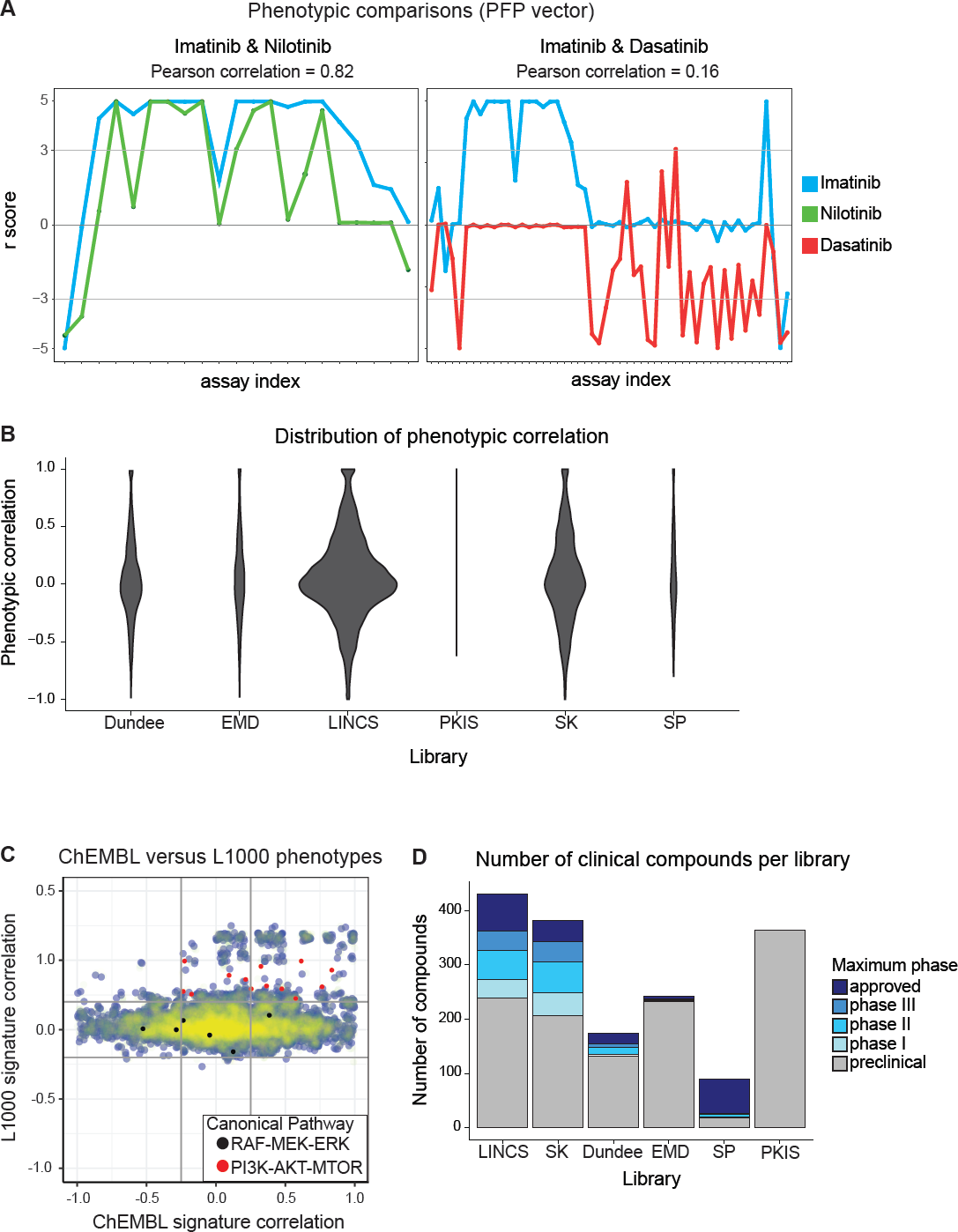
**Comparison of induced cellular phenotypes across kinase inhibitor libraries.** (A) Left panel: phenotypic fingerprint for Imatinib (yellow) and Nilotinib (green), which have a Pearson correlation of 0.82. Right panel: phenotypic fingerprint for Imatinib (yellow) and Dasatinib (red), which have a Pearson correlation of 0.16. Only assays for which either compound exhibited a robust z-score (r-score) greater than 3 or smaller than −3 are shown. (B) Distribution of pairwise correlations among phenotypic fingerprints across compounds in different libraries normalized to the number of compounds for which phenotypic information is available in ChEMBL. (C) Comparison of ChEMBL-derived PFP and L1000 Connectivity Map (C-MAP) correlations. Black and red dots highlight compounds targeting the PI3K-AKT-MTOR or RAF-MEK-ERK pathways (see text for details). (D) Number of compounds in various phases of clinical development for each library

We find that the LINCS and SK libraries have the lowest correlations when all pairs of PFP vectors are considered whereas the PKIS and SP libraries have the highest correlation and are therefore the least diverse (Figure 2B). A potential complication in this comparison is that the number of phenotypic assays reported in ChEMBL varies from compound to compound and also, on average, from library to library. Thus, libraries might appear to have lower phenotypic diversity simply because they are insufficiently annotated. This issue is particularly acute for compounds in the PKIS library, few of which are extensively annotated.

Gene expression profiles from cells exposed to small molecules provide an independent means of measuring a phenotype. A large number of such profiles (˜ 1.5 x 10^6^) is available through the Connectivity Map (CMAP; http://clue.io/cmap)^34,35^. For compounds in the LINCS library we compared correlation coefficients for pairs of PFP vectors with correlation in CMAP L1000 profiles (using level 5 characteristic direction vectors^36^). The great majority (73%) of compounds exhibited phenotypic correlation (set at a threshold of r ≥ 0.2 L1000, r ≥ 0.25 PFP) or non-correlation (|r| < 0.2 L1000, r < 0.25 PFP) by both metrics but a subset (27%) where discordant (Figure 2C). For example, when pairs of compounds targeting PI3K-MTOR-AKT signaling were compared, L1000 profiles were more likely to be correlated than PFP values, which is consistent with data showing that phenotypic responses to such drugs can be highly cell-type specific^36^. A similar phenomenon was observed for compounds targeting the RAF-MEK-ERK network.

The stage of clinical development is the final additional criterion on which to evaluate compounds in a library since the study of approved or investigational therapeutics has intrinsic value, even when such compounds do not substantially contribute to library diversity. LINCS has the greatest number of approved drugs (n=67; 14% of total); SK has the greatest number of compounds that are currently or have previously been tested in phased clinical trials (n=137; 32%); SP has the highest proportion of approved drugs (n=54; 57%) and PKIS the lowest (0%) (Figure 2D). Below we describe how clinical stage and compound diversity can be balanced against each other during library assembly.

### Compound selectivity and library coverage

Compounds are commonly described with respect to their nominal target (e.g. a “BRAF-inhibitor”, in the case of Dabrafenib). However, this does not mean that the nominal target is the highest affinity target or the most biologically or clinically relevant. Crizotinib, for example, was developed as an inhibitor of the cMET receptor, but was subsequently found to inhibit ROS1 and ALK kinases. In culture it was observed that apoptosis was induced specifically in NSCLC lines carrying an EML4-ALK translocation^37^ and the FDA initially approved the drug for this indication^38^. In general, it is difficult to determine which target is most relevant for a specific phenotype, nor is it always clear that any single target – rather than a collection of targets – is responsible for biological activity. We therefore used the *selectivity score* developed by Wang et al^7^, which assesses selectivity for each compound-target interaction, rather than for a compound overall, while accounting for data bias.

*Selectivity score* evaluates compound-target pairs based on the log-distribution of on-target affinities relative to the log-distribution of off-target affinities; this yields a *selectivity score* for every target against which a compound has been tested. *Selectivity score* measures the magnitude of difference in the first quartile of the on-target and off-target distributions (q_1_) and determines the rank sum *p-*value to assess significance. The *selectivity score* accounts for the fact that many compounds have only been evaluated on a relatively small number of targets, which can falsely give the impression that a compound is known to be highly selective. For example, selectivity is judged to be higher for compound ***A*** than compound ***B*** if ***A*** has been tested against a large number of targets and found to not bind most of them even if ***B*** has a higher affinity for a particular target of interest but has not be tested against alternative targets. It should be noted that like other features used to assess compounds in this study, *selectivity score* changes over time as additional data become available.

*Selectivity score* is a continuous variable but for convenience we also assigned descriptive labels to different classes of compound-target interaction. *Most Selective* (*MS*) interactions meet four criteria: (i) the difference in q_1_ values computed for the distributions of on- and off-target data is not less than 100-fold, (ii) the distributions of on-target and off-target binding affinities differ with *p-*value below 0.1 (iii) the compound has at least four-fold more off-target than on-target affinity measurement (so that data bias is below 20%), (iv) at least two published reports establish that the affinity for drug-target interaction is less than 100 nM. *Semi-Selective* (*SS*) interactions have slightly less stringent criteria: (i) the difference in q_1_ values is not less than 10-fold, (ii) the *p-*value for on and off-target binding affinities is below 0.1 (iii) at least four publication report affinity under 1 µM (because the affinity cut-off is less stringent than for interactions meeting *MS* criteria, more data-points are needed to establish confidence in binding). *Poly-Selective* (*PS*) interactions are those for which: (i) q_1_ values for on- and off-target binding are similar, (ii) on-target q_1_ is under 9 µM (iii) data bias is under 20%. *Unknown* (*UN*) interactions are those for which the compound has not been sufficiently probed for off-target binding but fulfills the *PS* requirements with respect potency and affinity for the nominal target.

The requirement for such a multi-parametric approach is exemplified by the tool compound PP121, whose nominal targets are BCR/ABL1 and PDGFR, and Dasatinib, an approved drug whose nominal targets are BCR/ABL1, Src family kinases and PDGFR. Dasatinib has more known targets with an affinity between 1nM and 1 µM than PP121 (Figure 3A; red circles), but it has also been tested on a substantially larger number of kinases. In many cases, it exhibits no inhibitory activity against these off-targets (K_i_ > 10 µM, black). Thus, the *selectivity score* for Dasatinib (score = 1.1) is substantially higher than that of PP121 (0.44). We found that the LINCS and SK libraries had the largest number of kinases with *MS* interactions and the PKIS library the fewest (Figure 3B).

**Figure 3.**
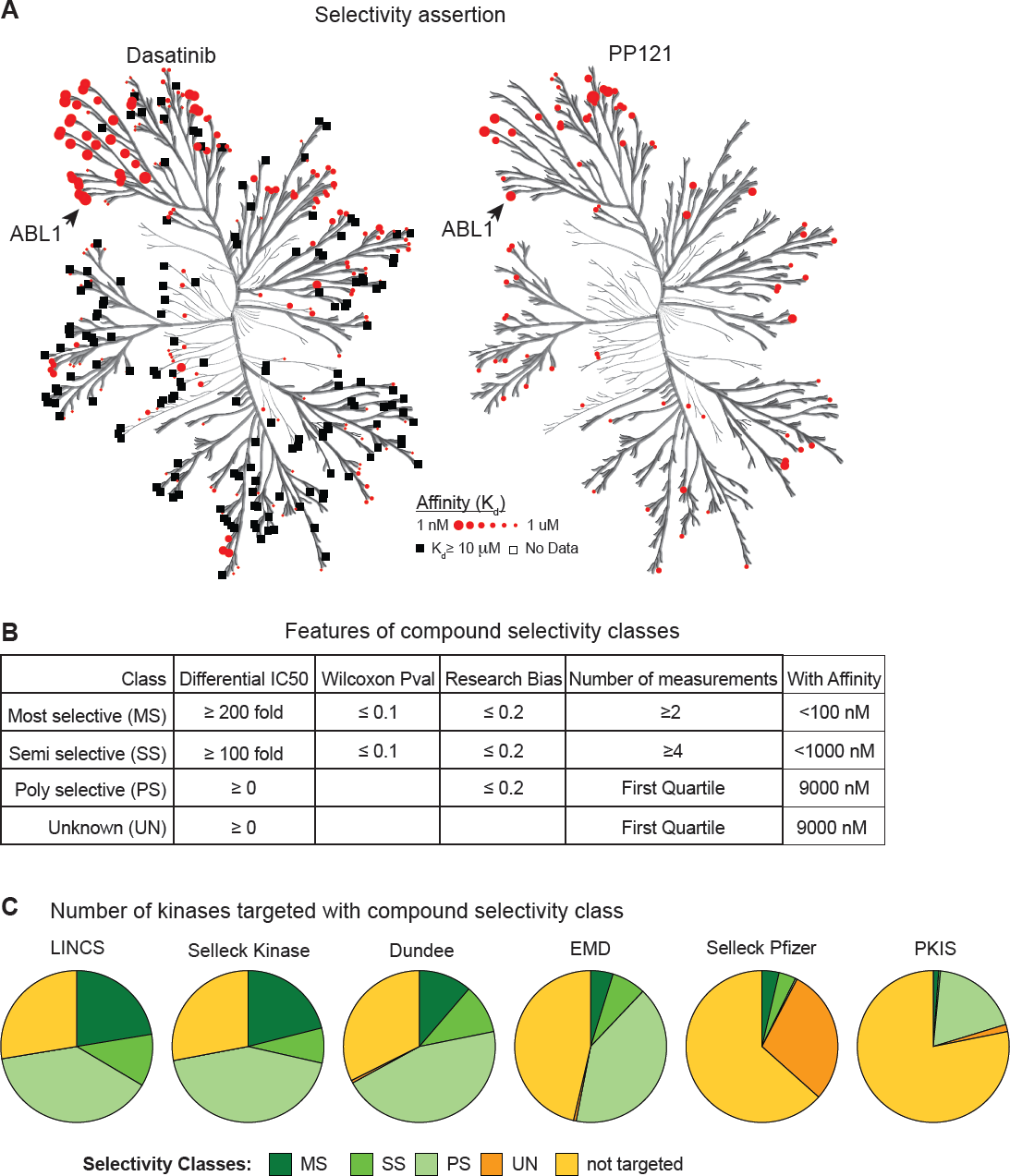
Comparisons of selectivity across libraries. (A) Illustrative example of selectivity assertions with incomplete target information. Estimated binding affinities for Dasatinib (right) and PP121 (left) are shown on a kinase tree. Red circles denote confirmed binding (bigger circles indicate higher affinity), black squares denote confirmed non-binding (K_d_ > 10uM), the absence of data point (no color) denotes an absence of information. (B) Features of compound selectivity classes. Note that poly-selective (PS) and unknown (UN) selectivity classes require that first quartile of all affinities measured be less than 9000 nM. (C) Pie charts show the number of kinases inhibited within a specified selectivity class per library. The highest selectivity achieved for each of the 545 human kinases is shown.

### Evaluating compound similarity on binding spectrum

To compare compounds based on all known targets and known non-targets, we sought a means to summarize all available data on a compound, including dose-response data, profiling data and curated annotations from the literature (e.g. “*drug X was found to not inhibit* …” or *“was equipotent for …”*). The resulting *Target Affinity Spectrum (TAS*) vectors summarize binding information from multiple assay formats weighted for the degree of evidence for high affinity binding and also for non-binding (K_d_ > 10 µM). TAS vectors contain some best-guess weighting parameters and are therefore less precise than *selectivity scores*, but they provide a more complete picture of available data (Figure 4A).

**Figure 4.**
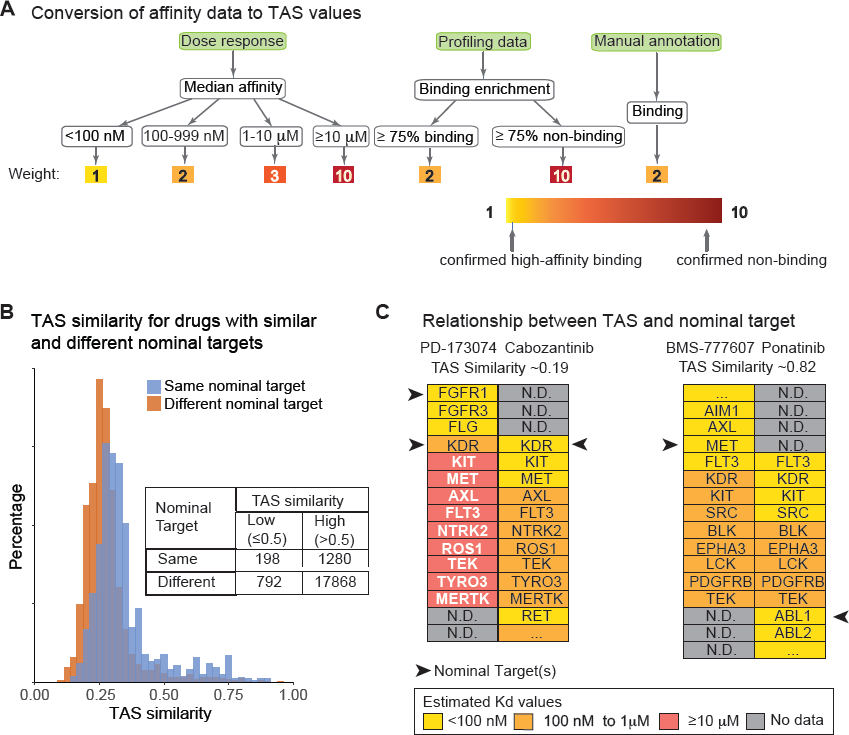
Comparisons of target affinity spectra across libraries. (A) Schematic illustrating the conversion of three types of data on target binding (dose response, one-point inhibition and manual annotation) into the binding assertions summarized by TAS vectors. Assertions of 1 (yellow) correspond to confirmed high-affinity binding. Assertions of 10 (red) correspond to confirmed non-binding. Target assertions were used to quantify similarities in target affinity spectra. (B) Distribution of target affinity similarities (calculated using weighted Jaccard similarities) for compounds having the same (blue) or different (brown) nominal targets. Inset table depicts the number of compounds used for the analysis. (D) Illustrative examples of the lack of correlation between a target spectrum and nominal drug targets. PD-173074 and Cabozantinib illustrate a pair of compounds with the very similar nominal targets but different target spectra. BMS-777607 and Ponatinib illustrate a pair with different nominal targets, but similar target spectra. Black arrows denote nominal target(s); targets are listed in alphabetical order by affinity class. As it happens, in the example shown neither compound had a target binding weight corresponding to an estimated K_d_ of 1–10 µM.

For the LINCS library, TAS vectors varied in length from a median of 10 features for probe compounds (range 1-394) and 15 for approved drugs (range 1-420), (Wilcoxon rank-sum test p=0.07). In no cases did a TAS include assertions for all 545 human kinases^39^, let alone for all ˜3000 members of the ligandable genome^40^, illustrating the sparseness of target annotation. TAS similarities for all compound pairs were calculated using a weighted Jaccard similarity metric (see methods). We found that TAS similarity was robust to at least 20% random variation in user-assigned weights (Figure S3).

We compared TAS vectors computed from ChEMBLV22_1 and other resources to data from a systematic assessment of kinase-inhibitor binding recently published by Klaeger et al^41^ that uses whole-cell lysates and mass spectrometry to profile compound-target binding (ChEMBL does not yet include this data). Surprisingly, the target coverage of the two datasets only partially overlapped, with a typical LINCS compound having one-third of its targets annotated in both datasets and one third in only one of the two datasets (Figure S5A). The reasons for differences in target spectrum as curated by ChEMBL and measured by Klaeger et al. are not yet clear and may relate to the range of targets evaluated in each case, but for targets annotated in both sources, 90% of the target-affinity assertions were congruent (Figure S5B).

We also asked whether knowledge of a molecules’ nominal target was informative about target spectrum by comparing TAS similarities for compounds with the same or different nominal targets. Remarkably, we found that the distribution of TAS similarities was very similar for compounds having the same nominal targets and for compounds having different nominal targets (Figure 4B). Thus, relatively little information about target spectrum can be discerned from nominal target alone. For example, consider two pairs of compounds: (i) PD-173074 and Cabozatinib and (ii) BMS-777607 and Ponatinib (Figure 4C). PD-173074 and Cabozantinib are structurally dissimilar (structural similarity 0.13) but overlap in their nominal targets, which are KDR and FGFR1 receptors for PD-173074 and KDR for Cabozantinib (black arrows in Figure 4C). These two compounds have non-overlapping secondary targets and thus, their TAS similarity is 0.19. In the case of BMS-777607 and Ponatinib (structural similarity 0.12), the nominal targets are different (MET and ABL1, respectively) but the secondary targets are very similar (Figure 4C right panel) and their TAS similarity is therefore 0.82. We propose that TAS, or a functional equivalent, rather than nominal target be much more widely used when analyzing the effects of compounds on cellular physiology.

### Relationship among structural, phenotypic and target similarity

For many compounds, data on target spectrum and induced cellular phenotype are sparse and *selectivity score* or TAS vectors might not be computable. We asked whether, in these cases, structural similarity could be used as a surrogate. In general, we found that structural similarity is a relatively poor predictor of phenotypic similarity: for example, the likelihood that compounds with structural similarity 0.7 (representing the Tanimoto similarity of Morgan2 fingerprints) or higher will induce similar phenotypes (i.e. exhibit a PFP correlation ≥ 0.9) is only 10% (Figure 5B). We also found that for compounds in the LINCS library (which has high structural diversity), structural similarity and target similarity were not correlated (Figure 5C). This was confirmed in the case of a subset in the PKIS library that shared the same core structure: structural similarity did not predict target similarity (Figure 5D). However, the converse appear to be true: when compounds had low structural similarity (≤ 0.2; Figure 5A) they were more likely to induce different phenotypes and have different target spectra. Thus, structural, phenotypic and target similarity contain non-redundant information about small molecule similarity. However, when data on target selectivity is missing, it is possible to exploit the finding that structures with very low similarity (≤ 0.2) have the highest likelihood of being diverse at the phenotypic and target levels.

**Figure 5.**
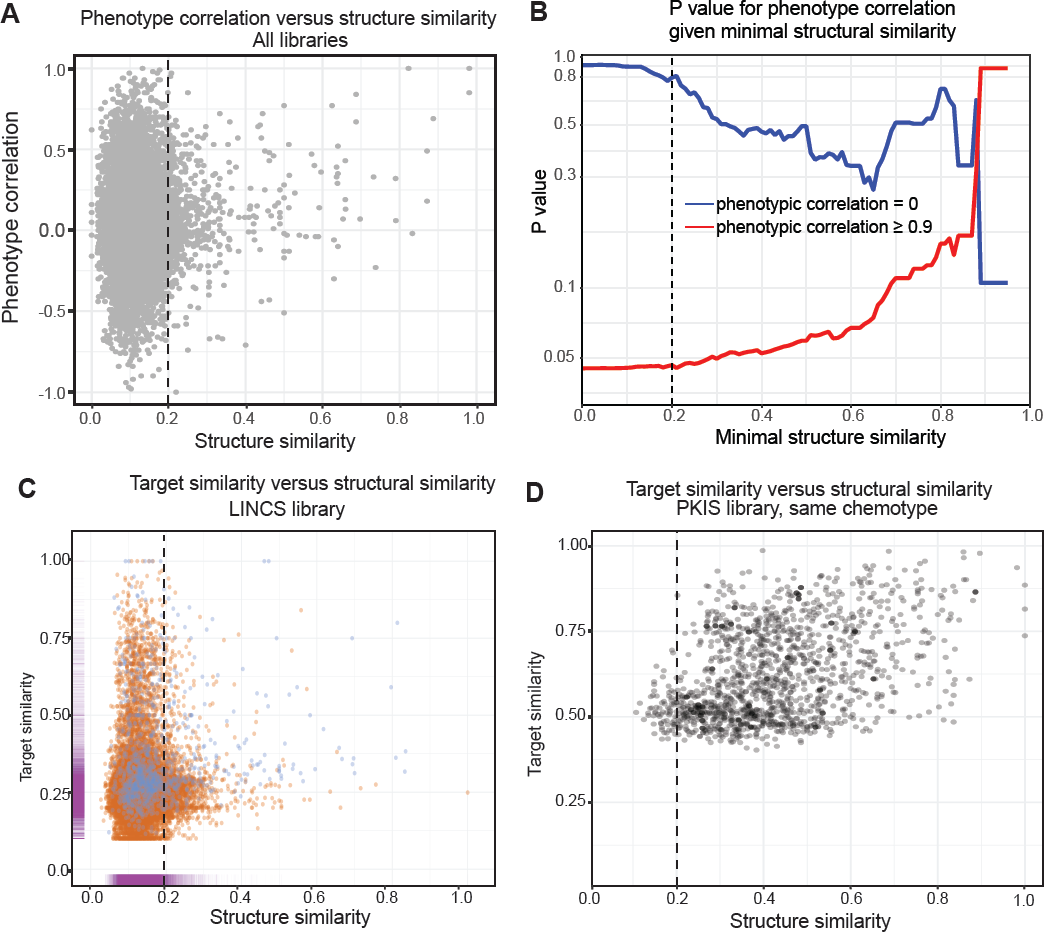
Comparison of structural, phenotypic and target similarity. (A) Comparison of phenotypic correlation and structural similarity for all libraries. Only a very small subset of the data is correlated. Dotted line denotes the threshold of 0.2 used for library design. (B) Conditional p-values for phenotypic-correlations given a specified minimal degree of structural similarity. (C) Comparison of target similarity and structural similarity for the LINCS library and (D) for the PKIS library In panel C, compound-pairs with same nominal target are plotted in blue, compound-pairs with different nominal targets are plotted in brown. The marginal distributions for all compound pairs are plotted in magenta on the axes.

### Designing an improved kinase inhibitor and a mechanism of action library

To enable construction of new libraries using the methods described above, we created the *LibraryR* software tool and deployed it as a web application using R-Shiny (http://www.smallmoleculesuite.org). For a user-specified set of genes *LibraryR* mines ChEMBL to create libraries containing (i) two compounds maximally specific for each gene of interest but having structural similarity ≤ 0.2, so that overlap in secondary targets is minimized (ii) all available compounds at a user-specified clinical stage (e.g. clinical phases 1-3 or approved) that also bind to the genes of interest with user-specified affinity (default Ki ≤ 1µM). *LibraryR* also calculates the *selectivity score* for each compound and assigns each compound to selectivity classes (*MS, SS, PS, UN;* Figure 6A). We used *LibraryR* to construct an improved version of the LINCS kinase library (the Laboratory of Systems Pharmacology *LSP-OptimalKinase* library; Table S14) and a mechanism of action library that optimally targets the liganded genome (*LSP-MoA*; Table S15).

**Figure 6.**
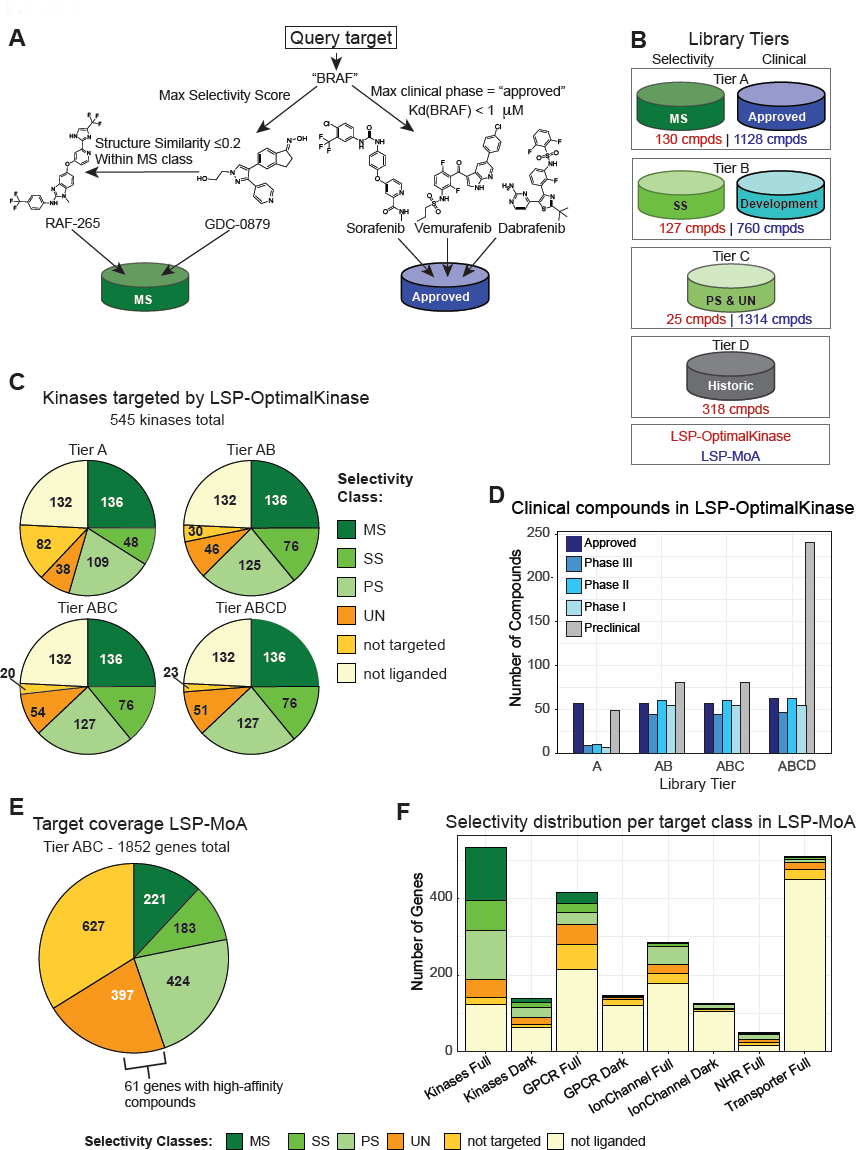
Creating a custom chemical genetics library. (A) Illustration of the compound selection procedure for a query target “BRAF” based on selectivity score and clinical phase. In this case, all compounds shown belong to Tier A as they either have *MS* selectivity or are clinically approved drugs. (B) Overview of the tiered design of the MoA and LSP-OptimalKinase libraries based on selectivity class and phase of clinical development. (C) Number of kinases inhibited at a specified selectivity class based for the LSP-OptimalKinase Library. The highest selectivity achieved for each of the 545 human kinases is shown. (D) Clinical grade compounds in the LSP-Optimal Kinase library by tier. (E) Target coverage by selectivity class for the liganded in the LSP-MoA library (F) Selectivity with which different target classes can be inhibited by the LSP- MoA library. “Full” full refers to the complete target class and “dark” to the understudied portion of the class as curated by the NIH IDG program (see text for details and Supplementary Tables 7, 10–13).

Both the *LSP-OptimalKinase* and *LSP-MoA* libraries are tiered based on selectivity and clinical phase to allow sub-libraries of different sizes to be assembled (Figure 6B). Tier A is the minimal library and contains only those compounds (i) binding the specified list of genes with *MS* selectivity and (ii) FDA approved drugs binding more strongly than the affinity cutoff. Tier B adds compounds with SS selectivity that target genes not covered by MS selectivity plus all compounds that bind the genes of interest and are in clinical development (clinical phase I-III). Tier C adds compounds from *PS* and *UN* specificity classes to maximally cover the user-specified list. Tier D includes all compounds historically included in a pre-existing library or present for other reasons (e.g. as positive and negative controls; Figure 6B). Inclusion of Tier D molecules is important to allow historical comparisons, gather phenotypic fingerprints and increase the chance for serendipitous discovery, which scales roughly with library size. Depending on the number of tiers selected, the LSP-OptimalKinase library ranges in size from 130 compounds to 590 compounds.

The use of *LibraryR* substantially increased target coverage and decreased the number of compounds in LSP-OptimalKinase relative to the LINCS Kinase Library we had assembled by hand several years ago. The 130 compounds in Tier A of *LSP-OptimalKinase* target 136 kinases with *MS* selectivity as compared to 117 kinases by the 495 compounds in the original LINCS library. Adding 127 Tier B compounds extends *MS* and *SS* coverage to 37 additional kinases (Figure 6E, Figure 3C) while including many more compounds in clinical development (Figure 6F). However, the further addition of 343 Tier C and D compounds extends coverage to only 10 additional kinases at *PS* and *UN* selectivity. Thus, computation-driven library design can reduce the size (and cost) of screening libraries and the studies that use them, while increasing target coverage and maintaining library diversity.

The LSP-MoA library is designed to target the 1852 members of the liganded genome (Table S9) about 12% of which can be bound with *MS* selectivity, 10% maximally with *SS* selectivity, 44% maximally with PS or UN selectivity; 34% of kinases are bound only by compounds that are known to have a higher affinity for another target (Figure 6C). In the case of targets bound with UN selectivity, 61 involved relatively tight binders (compounds with K_d_ < 100 nM) for which little selectivity data is available. Such compound-target pairs would benefit disproportionately from more data on possible off-targets (e.g. obtained by profiling). We analyzed LSP-MoA for its coverage of five therapeutic target classes: kinases, G-protein-coupled receptors (GPCRs), ion channels (ICs), nuclear hormone receptors (NHRs), and transporters (supplementary Tables 7, 10-13). Kinases were by far the best covered with 136 of 545 kinases (25%) at *MS* selectivity, followed by 28 of 416 GPCRs (7%), 2 of 49 NHRs (4%), 2 of 285 ICs (0.7%) and 1 of 510 transporters (0.1%; Figure 6D). The same trend was also observed for the understudied (‘dark’) ligandable genome as recently curated by the NIH (https://commonfund.nih.gov/idg Tables S7, S10, S11).

## Discussion

The ideal focused library of annotated, biologically active small molecules comprises the minimal number of compounds necessary to cover a gene set of interest selectively while optionally including additional approved and investigational drugs. Achieving optimal coverage is difficult in practice since many small molecules bind multiple targets and data on target spectra are nearly always incomplete and subject to poorly understood experimental confounders. Moreover, in many cell-based screening projects, the focus is not on targets *per se* but instead on induced cellular phenotypes. In this paper, we develop a set of criteria for evaluating and constructing small molecule libraries based on binding selectivity, target coverage and cell-based phenotypes as well as chemical structure and stage of clinical development. We find that data on structure, target spectrum and induced phenotypes are non-redundant, demonstrating that compounds should only be labeled as being functionally ‘similar’ or ‘different’ when evaluated on diverse types of data.

We have codified our approach to scoring and assembling compound collections in three web-based applications: *SelectivitySelectR, SimilaritySelectR* and *LibraryR*, all of which draw data from ChEMBL and other user-specified sources. For a given list of targets, libraries are constructed so that each target is ideally bound by two high-affinity compounds with diverse target spectra and induced cellular phenotypes. Such data are incomplete but it is possible to increase the likelihood that a library will be functionally diverse by including compounds with non-similar structures – defined in this study as a Tanimoto similarity of Morgan2 fingerprints ≤ 0.2. This finding is in agreement with the general consensus^18,42^ although our similarity cutoff is more stringent than usual. Additional therapeutics, beyond two per target, are included in subsequent tiers of the library because such compounds have intrinsic value for translational studies and drug repurposing. The bottom tiers in a library attempt to cover additional targets at lower levels of selectivity and also include historical compounds needed for comparison to previous libraries.

When six widely available kinase inhibitor libraries were evaluated, we found that the LINCS and SK libraries had the greatest diversity, broadest coverage of the kinome and the largest number of approved and investigational drugs. These were also the largest libraries analyzed, which increases the cost of creating and using them, particularly when complex dose-response or phenotypic assays are involved. The Dundee, EMD and SP libraries were substantially smaller (19% – 54% the size of LINCS), but underperformed in one or more metrics. The PKIS library scored poorly with respect to structural and phenotypic diversity and number of approved drugs, but did have the largest number of compounds not included in any other collection. When we used *LibraryR* to design a new *LSP-OptimalKinase* library we found that a collection of 256-compounds (Tiers A plus B; see Figure 6) outperformed all existing kinase libraries with respect to selectivity, target coverage, structural diversity and number of approved and investigational drugs. This library is also smaller than all but the existing SP library, which has substantially poorer coverage of the kinome.

Because our approach is data-driven and considers only public information, we have observed some differences between our top picks and first choice compounds suggested by others^18–20^. For example, www.chemicalprobes.org lists BI-D1870 as a selective inhibitor of the RPS6KA1 kinase, while we score this drug-target pair as being of *unknown selectivity*. In the published literature D1870 has only been tested against three targets, which *LibraryR* considers insufficient evidence of selectivity. However, *LibraryR* has an option to add compounds from expert-opinion lists and then rank them relative all other compounds in a collection.

We also generated a list for a 3202-compound “mechanism of action library (LSP-MoA) that optimally covers the liganded genome and should be of general utility in semi-focused screening campaigns. We found that kinases were the target class best covered by this library, GPCRs were a distant second and many other multi-protein families had few if any selective inhibitors. This emphasizes the relative sparsity of selective coverage of targets considered ‘ligandable,’ at least by compounds whose structures are in the public domain. However, our approach to analyzing and designing compound libraries is data-driven so that the MOA library can be steadily improved as new data become available.

We expect the methods described here to assist in a variety of informatic, chemical biology and drug repurposing studies. For cell-based experiments, our tools can help to identify multiple, selective, and structurally diverse compounds against one or more target(s) of interest, thereby reducing the likelihood that off-target effects go unrecognized. Machine learning applications should also be able to make use of *target affinity spectrum* to investigate molecular profiling data on drug-treated cells, particularly when multiple compounds with the same or similar nominal targets are compared. It should also be possible to further improve the use of large-scale profiling and phenotypic data (e.g. the Connectivity Map^35,43^ and Cancer Cell Line Encyclopedia^44^) to optimize compound collections. Finally, we hope that the ready availability of data on the full spectrum of compound activities will replace nominal target alone as a means to describe mechanism of action.

## >Acknowledgements

This work was supported by NIH grants U54-HL127365 (NM, PKS and NG), U24-DK116204 (NM, PKS) U54-HL127624 (NAC and MM) and by the Novartis Institutes for BioMedical Research (YW, EL and JJ). We thank V. Becker for creating the literature annotation table for compounds in LINCS.

## Methods

**Data and resource availability**

| File name | File type | Functionality |
| --- | --- | --- |
| 01_chemical_identity&similarity.py | Source code | Match identity of chemical structure |
| 02_collecting_data_chembl.r | Source code | Collect data from ChEMBL |
| 03_calculating_PFPsimilarity.r | Source code | Calculate the phenotypic fingerprint correlation between compound pairs |
| 04_import_gctx.py | Source code | Import L1000 data and calculate correlation between compound pairs |
| 05_calculating_selectivity.r | Source code | Calculates *Selectivity score* |
| Suppl_Table02_compoundmap.csv | Data table | Maps compound identifiers from sources used in this manuscript |
| Suppl_Table03_target_map | Data table | Maps target identifiers from sources used in this manuscript |
| Suppl_Table04_curated_literature_annotations_LINCS.csv | Data table | Contains the manually curated target information from literature |
| Suppl_Table05_onepoint_kinomescan_vs_Kd.csv | Data table | Compares DiscoverX KinomeScan read-outs to dose-response Kd values |
| Suppl_Table05_onepoint_Dundeescan_vs_Kd.csv | Data table | Compares Dundee profiling read-outs to dose-response Kd values |
| Suppl_Table07_kinase_table.csv | Data table | Lists human kinases and their participation in 'dark' and 'liganded' genome |
| Suppl_Table08_TAS_vectors_LINCS_cmpds.csv | Data table | Contains the TAS vectors for compounds in LINCS |
| Suppl_Table09_liganded_genome_v22_1_20171217.csv | Data table | Lists the liganded genome on Dec 17<sup>th</sup> 2017 |
| Suppl_Table10_gpcr_table.csv | Data table | Lists human GPCRs and their participation in 'dark' and 'liganded' genome |
| Suppl_Table11_ionchannel_table.csv | Data table | Lists human Ion channels and their participation in 'dark' and 'liganded' genome |
| Suppl_Table12_NHR_table.csv | Data table | Lists human nuclear hormone receptors and their participation in the 'liganded' genome |
| Suppl_Table13_transporter_table.csv | Data table | Lists human transporters and their participation in the 'liganded' genome |
| Suppl_Table14_cmpdlist_LSP_OptimalKinase.csv | Data table | Compounds in the LSP-OptimalKinase library with mentioning of reason included |
| Suppl_Table15_cmpdlist_LSP_MoA | Data table | Compounds in the LSP-MoA library with mentioning of reason included |
| Table note: the files described above are available in supplementary material. |  |  |

All source code can be found online on www.github.com/sorgerlab/smallmoleculesuite

All source code can be found online on www.github.com/sorgerlab/smallmoleculesuite

### Compute chemical identity and similarity

Compounds from the different libraries were matched to each other and their ChEMBL equivalent based Tanimoto similarity of their Morgan2 fingerprints. Those compounds that had multiple matches with 100% similarity were curated manually. See script “01_chemical_identity&similarity.py”

### Collecting data from ChEMBL V22_1

A local copy of ChEMBL V22_1 was installed. Both phenotypic and target data were collected from the ACTIVITIES table. The two datatypes were distinguished based on assay annotation detailed in script “02_collecting_data_chembl.r”. Target identifiers were converted to ENTREZ gene identifiers using ‘gene_symbols’ from the COMPONENT_SYNONYMS table and targets with multiple or no matches were manually checked. See Table S3A-3C for mapping. Maximum clinical development stage was obtained from the MOLECULE_DICTIONARY table.

### Calculating phenotypic fingerprint (PFP) correlation

Results from qualifying phenotypic screens were normalized to robust z-scores (r-scores). For each compound pair, Pearson’s correlation was calculated on the subset of assays in which both compounds had been measured and least one of the compounds was active (i.e. r-score ≥3 or r-score ≤-3); correlation for pairs that had less than five of such assays were excluded. See script “03_calulating_PFPsimilarity.r”.

### Calculating L1000 correlation

Level 5 of the L1000 data was downloaded from GEO accession number GSE92742 on May 1^st^, 2017 and filtered to retain only compounds in the LINCS library. For each pair of compounds, all Pearson correlations were calculated between perturbations measured in the same cell line, at the same time point, and with same treatment concentrations (see script ‘04_import_gctx_allconc.py’). The final value reported in Figure 2C is the median distance for each pair of drugs.

### Calculating *selectivity score*for compound-target interactions

The *selectivity score* for compound-target interactions was calculated according to an adapted version of Wang et al^7^, so that:

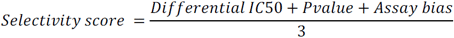

with:

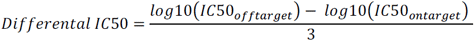

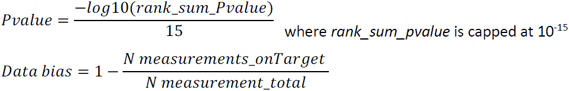

Selectivity classes, based on differential IC50, Pvalue and Data bias, were determined as outlined in Figure 3B. See script “05_calulating_selectivity.r”.

### Calculating Target Affinity Spectrum (TAS) similarity

The assignment of weights in the TAS vector was customized to each assay-type. In general, a weight of 1 denotes the highest confidence in binding and a weight of 10 denotes the highest confidence in confirmed non-binding. Dose response data was obtained from ChEMBL, DiscoverX and Kinative kinome profiles. Binding values were given a weight 1 if K_d_ <100nM, a weight 2 if 100nM ≤ K_d_ <1µM, a weight 3 if 1µM ≤ K_d_ <10µM, and weight 10 if K_d_ ≥10 µM. Annotations of nominal target were obtained from manual literature searches) and was assigned a weight of 2. One-dose data from profiling methods was obtained from weighted based on the retrospective probability of binding as explained in text.

Five different data sources were used to compose the TAS: (i) dose-response data from Chembl v22_1 (see above), (ii) dose-response data from DiscoverX kinomescan, (iii) profiling data from DiscoverX at doses 0.1uM; 1uM; 10 uM, (iv) profiling data from Dundee (http://www.kinase-screen.mrc.ac.uk/kinase-inhibitors) at doses 0.01uM; 0.1uM; 1uM; 10uM, (v) curated nominal target annotations. Targets were normalized to gene_ids using gene symbol matches and manual conversions. Binding enrichments for datasources iii and iv were calculated per concentration per source see Figure S2, Figure S2, and Table S4 for conversion results. Nominal targets were extracted from manual searches in pubmed and by mining vendor information. Binding assertions were assigned as illustrated in figure 4A. TAS vectors of compound pairs were joined and entries where one of the compounds had an assertion ≤3 were kept. If the joined vector had a length ≥5, a weighted Jaccard similarity coefficient was calculated with the formula:

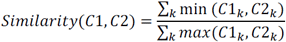

see Table S8 for full TAS vectors.

### Assembling LSP-OptimalKinase and LSP-MoA libraries

LSP-OptimalKinase was developed to target human kinases (Table S7), LSP-MoA was developed to target the liganded genome (Table S9). In the selectivity arm, the compound with the highest selectivity score was selected for each target. Then, a second compound within that same selectivity class (MS, SS, PS, UN) and a structural similarity ≤0.2 was selected. For targets with no second compound in MS class with sufficient structural dissimilarity, the SS class compounds were sourced to supply the second compound. The clinical development arm was assembled by selecting all molecules with max_phase 1-4 in MOLECULE_DICTIONARY table that had an affinity <1000 nM for one targets aimed at. Tiers were developed as illustrated in figure 5b. The members of molecular target classes for which the LSP-MoA library was analyzed were obtained from http://www.guidetopharmacology.org/download.jsp (accessed Dec 15th, 2017).

### The Small Molecule Suite graphical user interface

The small molecule suite comprises the applications *SelectivitySelectR*, *SimilaritySelectR* and *LibraryR*. For *SelectivitySelectR*, the *selectivity score* and mean affinity were calculated for all small molecules in the LINCS compound collection for each target their affinity was measured. In the application, users can select a target of interest and see the mean affinity and selectivity for all LINCS compounds with a known affinity for this target. All results can be downloaded for further used. For *SimilaritySelectR*, the structural similarity, TAS similarity and PFP similarity was calculated for all compound pairs in LINCS. In the application, users can select a reference molecule of interest to identify molecules that are similar or dissimilar in structure, target spectrum or induced phenotype. The application allows users to download information on the reference compound plus three additional compounds. For *LibraryR*, compounds were selected based on selectivity, clinical development stage and expert opinion status. To enable selection for compounds on selectivity the *selectivity score* was calculated for all compounds-target pairs in ChEMBL V22_1 for which the target was a member of the liganded genome and the compound was known to bind at least one target in the liganded genome with K_i_ ≤ 10µM. For each target, the compound with the highest *selectivity score* and another compound with a structural similarity ≤0.2 within the same selectivity class (*MS, SS, PS* or *UN*). To select compounds on clinical development stage, all compounds with K_i_ ≤ 10µM against any target in the liganded genome in development phase I-IV were selected. Users can set stricter requirement for affinity when composing the library (default K_I_ ≤ 1µM). Compounds curated by expert opinion were obtained from www.chemicalprobes.org^18^ (only those rated with 4/4 stars) and a recent publication by Wang & Gray^19,20^.

